# Riluzole treatment paradoxically increases motoneuron excitability in ALS due to hyperactive homeostasis

**DOI:** 10.64898/2026.03.23.713695

**Authors:** Amr A. Mahrous, Bradley S. Heit, CJ Heckman

**Affiliations:** Department of Neuroscience, Feinberg School of Medicine, Northwestern University, Chicago, IL, USA, 60611; Department of Physical Therapy and Human Movement Sciences, Feinberg School of Medicine, Northwestern University, Chicago, IL, USA, 60611; Department of Physical Medicine and Rehabilitation, Feinberg School of Medicine, Northwestern University, Chicago, IL, USA, 60611; Shirley Ryan AbilityLab, Chicago, IL, USA, 60611

**Keywords:** Amyotrophic Lateral Sclerosis (ALS), Riluzole, Motoneuron Excitability, Hypervigilant Homeostasis

## Abstract

Riluzole is the most commonly prescribed among the limited approved therapies for amyotrophic lateral sclerosis (ALS), a neurodegenerative disorder characterized by progressive motoneuron loss and paralysis. It is thought to act by suppressing motoneuron excitability and glutamate release, but its clinical benefits are modest and often diminish over time. We previously showed that homeostatic mechanisms in the SOD1^G93A^ (mSOD1) mouse model of ALS are hyperactive and prone to overcompensation. Here, we tested whether such dysregulated homeostasis antagonizes the effects of riluzole. Wild-type (WT) and presymptomatic mSOD1 mice received therapeutic doses of riluzole in drinking water for 10 days, with untreated littermates of both genotypes serving as controls. Motoneuron excitability and synaptic inputs were then examined using intracellular recordings from the isolated sacral spinal cord. The data showed that chronic riluzole treatment increased motoneuron excitability and polysynaptic inputs in mSOD1 mice but produced no detectable changes in WT motoneurons. These results suggest that hyperactive homeostatic mechanisms in ALS counteract the suppressive effects of riluzole. Notably, mSOD1 motoneurons exhibited larger membrane capacitance than WT, consistent with their increased cell size at this disease stage. Riluzole treatment reduced motoneuron membrane capacitance in mSOD1 mice to the range observed in WT animals, indicating normalization of cell size and potentially reduction in metabolic demand. Together, these findings help explain the limited clinical efficacy of riluzole while revealing a previously unrecognized neuroprotective mechanism of the drug in ALS.

## INTRODUCTION

Riluzole is an FDA-approved therapy used to slow disease progression in amyotrophic lateral sclerosis (ALS) [1-3]. At the cellular level, riluzole blocks voltage-gated Na^+^ channels, including those that generate the persistent inward Na^+^ current [4-7]. Its neuroprotective effects are believed to arise from reducing presynaptic glutamate release and postsynaptic neuronal excitability [4, 5, 8], thereby mitigating excitotoxic stress [9, 10]. Despite these actions, riluzole’s clinical benefit is limited, providing modest extension of survival in ALS patients [3, 11]. In this study, we test the hypothesis that therapeutic effects of riluzole might be counteracted by homeostatic adaptations that emerge during prolonged treatment.

Homeostasis refers to mechanisms by which cells maintain stable internal conditions despite external perturbations. These mechanisms operate as negative feedback systems that restore physiological balance. Mitchell and colleagues proposed that ALS motoneurons may exhibit *hypervigilant* homeostasis, an excessively high-gain feedback state [12, 13]. Their conclusion was based on a meta-analysis of multiple studies examining motoneuron properties in the SOD1^G93A^ (mSOD1) mouse model of ALS across disease stages. The analysis revealed oscillatory patterns (alternating increases and decreases) in mitochondrial function, oxidative stress markers, and intracellular Ca^2+^ levels throughout disease progression [13]. Consistent with these findings, our laboratory and others have reported temporal oscillations in motoneuron size and excitability in the same mouse model [14, 15]. Because such oscillations are characteristic of an overactive feedback system, these observations support the idea that motoneuron homeostasis becomes hyperactive in ALS causing overcompensation [16].

It remains unclear whether prolonged treatment with excitability-suppressing drugs, such as riluzole, may lose effectiveness over time due to heightened homeostatic responses in ALS motoneurons. Interestingly, a clinical longitudinal study has shown that riluzole produces temporary reduction in cortical and peripheral excitability in ALS patients, with some parameters returning to baseline after several weeks of continued treatment [17]. To investigate the role of dysregulated homeostasis in this phenomenon, we examined the impact of extended riluzole treatment on motoneuron excitability in the mSOD1 mouse model. We focused on a presymptomatic disease stage when motoneuron abnormalities are present but significant degeneration has not yet occurred. Both WT and mSOD1 mice received clinically relevant doses of riluzole for ten consecutive days, after which spinal cords were harvested for electrophysiological measurements of motoneuron excitability and their excitatory synaptic inputs.

Our findings reveal that this treatment regimen induced alterations in the properties and behavior of mSOD1 motoneurons, but not WT motoneurons, supporting the concept of dysregulated homeostasis in ALS. The intrinsic excitability of mSOD1 motoneurons was increased following extended riluzole treatment, and their excitatory synaptic inputs were slightly enhanced. These findings suggest that riluzole’s effects are indeed counteracted by homeostatic adaptations. This challenges the prevailing view of the mechanisms through which riluzole exerts its therapeutic effects in ALS. In addition, the data revealed other riluzole-induced changes in ALS motoneurons that might be neuroprotective, uncovering previously unrecognized benefits of riluzole treatment. This study thus provides novel insights into ALS disease mechanisms and highlights new potential therapeutic targets.

## MATERIALS AND METHODS

### Ethical Approval

All procedures in this study were conducted in accordance with the National Institutes of Health *Guide for the Care and Use of Laboratory Animals* and were reviewed and approved by the Northwestern University Animal Care and Use Committee. Careful measures were taken to minimize distress experienced by the animals in this study.

### Animals

Mice used in this study were obtained from The Jackson Laboratory (Bar Harbor, ME, USA). Young adult male mice were utilized and consisted of either transgenic animals overexpressing the mutant human SOD1 gene (mSOD1) or wild-type (WT) controls. WT controls were B6SJLF1/J mice (Stock No. 100012, JAX). The mSOD1 mice were hemizygous for the high-copy mutant transgene B6SJL-Tg (SOD1*G93A)1Gur/J (Stock No. 002726, JAX), which expresses the human SOD1 gene carrying the G93A mutation and develops a rapidly progressive ALS phenotype.

### Experimental design

Both mSOD1 and WT mice received either water containing riluzole hydrochloride (100 µg/mL) or regular drinking water (control groups). Treatment was initiated at approximately 37 days of age, corresponding to pre-symptomatic stage, and continued for 10 days (Fig. 1A–B). Animals had ad libitum access to water throughout the treatment period. Based on cage-level measurements of water consumption and average body weight (Fig. 1C), the estimated riluzole dose was approximately 22 mg/kg/day. This dose and method of administration (added to food or water) have been tested in mSOD1 mice and showed therapeutic benefits [1].

**Fig. 1:**
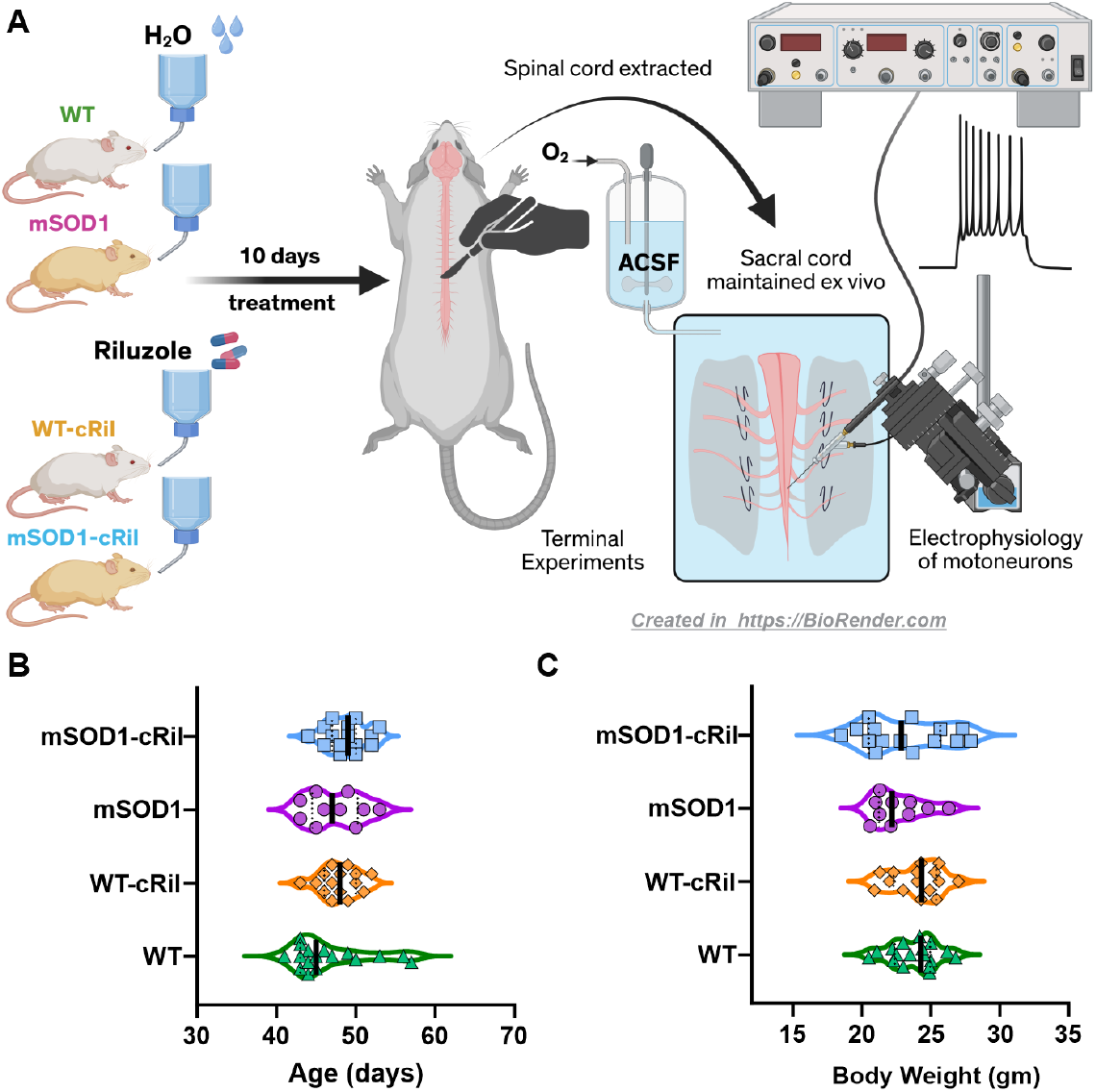
Experimental design. **A:** Young adult WT and mSOD1 mice either received riluzole in drinking water (100 µg/mL) or regular non-medicated water for 10 days. Following treatment, the whole-tissue sacral spinal cord was extracted in terminal experiments and maintained ex vivo. Electrophysiological recordings of motoneurons and ventral roots were then performed to assess changes in excitability and synaptic inputs. **B-C:** The age and body weight of the animals on the day of terminal experiments. There was no statistical difference in either of these parameters between experimental groups. Data was analyzed using Kruskal-Wallis test and Dunn’s multiple comparison test. N= 17 WT untreated, 15 WT-cRil, 10 mSOD1, and 15 mSOD1-cRil mice.

At the end of the treatment period, animals underwent a terminal experimental procedure. The sacral spinal cord was surgically isolated, maintained ex vivo, and subsequently used to assess motoneuron excitability and synaptic inputs.

### *Ex vivo* spinal cord preparation

Surgical isolation of the sacral spinal cord was performed as previously described [18, 19]. In brief, animals were deeply anesthetized with an intraperitoneal injection of urethane (≥ 2 g/kg body weight) and supplied with carbogen (95% O_2_, 5% CO_2_) through a face mask. A dorsal laminectomy was performed to expose the spinal cord, and the dura mater was opened longitudinally. The animal was then humanely euthanized by decapitation followed by transecting the cord at the lumbosacral enlargement (L5–L6). The whole sacrocaudal cord (segments S1 to Co2), with roots preserved, was maintained ex vivo as whole-tissue preparation.

The isolated cord was positioned ventral side up in a custom-built recording chamber and continuously perfused at room temperature (∼21°C) with oxygenated artificial cerebrospinal fluid (ACSF). The ACSF contained (in mM): 128 NaCl, 3 KCl, 1.5 MgSO_4_, 1 NaH_2_PO_4_, 2.5 CaCl_2_, 22 NaHCO_3_, and 12 glucose. Dorsal and ventral roots (DRs and VRs) were mounted on bipolar wire electrodes and sealed with petroleum jelly to prevent drying. Before recordings commenced, the preparation was perfused with ACSF for approximately 1 hour to allow physiological stabilization and to washout any residual riluzole.

### Electrophysiological recordings

Intracellular recordings from single motoneurons in the isolated sacral spinal cord were obtained using sharp glass microelectrodes, as described previously [19, 20]. Electrodes were filled with a 2 M potassium solution (1 M potassium acetate and 1 M KCl) and had tip resistances ranging from 35 to 55 MΩ. They were advanced into the tissue with a motorized micro-stepper (Kopf Instruments, CA). Motoneurons were identified by antidromic activation elicited through ventral root stimulation. Cells were accepted for analysis if the resting membrane potential was more negative than −60 mV and the antidromic action potential amplitude was 60 mV or larger. Recordings were performed using an Axoclamp-2B amplifier (Molecular Devices, CA) operating in either discontinuous current-clamp (DCC) or single-electrode voltage-clamp (SEVC) mode at switching rates that involved more than 20 cycles/membrane time constant as recommended by Manuel [21]). For ventral root (VR) recordings, roots were positioned on bipolar wire electrodes and connected to differential amplifiers (WPI, FL, USA) set to 1000× gain with a band-pass filter of 300 Hz to 3 kHz. Similarly, dorsal roots (DRs) were mounted on bipolar electrodes connected to a Grass stimulator (A-M Systems) to evoke synaptic inputs. DR stimulation consisted of bipolar pulses (0.1 ms) delivered at ≤ 0.066 Hz. Stimulation intensity was expressed as multiples of threshold (×T), where threshold was defined as the minimal current required to evoke the smallest detectable VR response. Both intracellular and extracellular signals were digitized at 10–20 kHz using a Micro3-1401 acquisition interface (CED, UK). Data were collected with Spike2® software (version 8–10, CED, UK) and stored for offline analysis.

### Data analysis

Motoneuron excitability was evaluated by eliciting repetitive firing using either slow triangular current ramps (0.5-1 nA/s for both ascending and descending phases) or square current pulses to construct frequency–current (F–I) relationships. The slope (gain) of the linear portion of the F–I curve was calculated using linear regression and served as a quantitative measure of intrinsic excitability. In addition to gain, the overall shape of the F–I relationship obtained during ascending and descending ramps was used to categorize motoneurons into four groups reflecting progressively greater excitability. Type I motoneurons exhibited firing rate adaptation, characterized by a clockwise F–I relationship and absence of sustained firing during the descending ramp. Type II motoneurons displayed a linear F–I relationship with overlapping firing frequencies during the ascending and descending phases. Type III motoneurons also showed a largely linear and overlapping F–I relationship; however, they exhibited sustained firing (a “tail”) during the descending ramp. Type IV motoneurons demonstrated higher firing frequencies on the descending ramp than on the ascending ramp, resulting in a counterclockwise F–I relationship, indicative of enhanced intrinsic excitability.

Synaptic transmission was evaluated by recording responses in ventral roots (VRs) and individual motoneurons following stimulation of the dorsal roots (DRs). For VR recordings, synaptic output was quantified as the peak-to-peak amplitude of the compound action potential (CAP). The ipsilateral monosynaptic CAP exhibited a short latency of approximately 2–2.5 ms, whereas the contralateral response showed a longer latency of 4–6 ms, consistent with additional synaptic delay [19]. Each CAP measurement represented the average of five consecutive trials. To account for variability between preparations, all ipsilateral and contralateral CAP amplitudes from a given VR were expressed as percentage of the root’s maximal response. The VR maximum response was defined as CAP amplitude evoked by ipsilateral DR stimulation at 10× threshold. VRs with a maximal peak-to-peak amplitude of less than 1 mV were deemed unhealthy and excluded from analysis.

In addition to VR recordings, excitatory postsynaptic potentials (EPSPs) were recorded intracellularly from single motoneurons in response to contralateral DR stimulation. For each cell, EPSPs were averaged across 3–5 trials, and quantified as the area under the membrane potential curve within a 0– 300 ms time window following stimulation.

Intrinsic electrical properties of individual motoneurons were assessed using intracellular recordings. Input resistance was determined from the slope of the current–voltage (I–V) relationship generated by injecting a series of hyperpolarizing current steps (500 ms duration, 0.25–1 nA). Voltage responses were measured both at the initial peak deflection (R_peak_) and at the steady-state plateau (R_plateau_). Input conductance was calculated as the reciprocal of input resistance. The sag ratio was calculated as the ratio of the peak voltage deflection to the steady-state (plateau) response. Rheobase was defined as the minimal amplitude of a depolarizing current pulse (50 ms duration) required to evoke an action potential. After determining rheobase, a brief, higher-amplitude current pulse (0.5 ms, approximately 10× rheobase) was delivered to reliably elicit a single spike for measurement of the medium afterhyperpolarization (mAHP). Because mAHP amplitude varies with membrane potential, it was measured at several resting membrane potentials. A linear regression of mAHP amplitude versus membrane potential was used to estimate the mAHP amplitude at −70 mV. This relationship was also used to calculate the conductance and reversal potential of the AHP, as previously described [22]. The half-decay time of the mAHP was defined as the interval from the peak amplitude to the point at which it decayed by 50% toward resting potential. We also measured persistent inward currents (PICs) using single-electrode voltage-clamp (SEVC) mode. Cells were held at −70 mV, and slow depolarizing voltage ramps (5 mV/s, 40 mV amplitude) were applied. After subtracting the linear leak current, the PIC amplitude was quantified as the peak inward current observed during the ascending phase of the voltage ramp.

The membrane time constant (τ_m_) was determined from the voltage response to brief, small-amplitude hyperpolarizing current pulses (−1 nA, 1 ms). Following current injection, the decay of the membrane potential was analyzed. The later portion of this decay, which reflects predominantly the passive membrane properties, was plotted on a semilogarithmic scale. A linear fit was applied to this region, and the membrane time constant (τ_m_) was calculated from the slope of this line. However, the voltage relaxation does not follow a single exponential time course in neurons with extended dendritic trees. Instead, it reflects multiple exponential components arising from current redistribution between the soma and dendrites. To separate these components, we used the method introduced by Rall [23]. Briefly, after determining τ_m_ from the slowest (dominant) exponential component, this component was mathematically subtracted (“peeled off”) from the original voltage trace. The remaining earlier, faster component was then isolated and fitted separately to obtain the first equalizing time constant (τ_1_). This peeling procedure is necessary because the membrane response of a spatially extended neuron cannot be accurately described by a single exponential decay. The slow component (τ_m_) primarily reflects the overall membrane charging time, whereas the faster equalizing component (τ_1_) reflects the redistribution of charge between the soma and dendrites. By separating these components, it becomes possible to estimate the electrotonic structure and total capacitance of the motoneuron based on cable theory. Assuming the motoneuron (soma and dendrites) can be approximated as an equivalent cylinder with uniform membrane resistivity and capacitance, the electrotonic length (L) was calculated from T_m_ and T_1_ using the relation: L = π / √(T_m_ / T_1_ − 1). The total membrane capacitance (C) was then estimated according to: C = (τ_m_ × L) / (R_peak_ × tanh(L)) [23-25].

#### Drugs and chemicals

Riluzole was purchased from HelloBio (Princeton, NJ), in the form of riluzole hydrochloride (water-soluble). All chemicals used to prepare the physiological solutions were purchased from Sigma-Aldrich^®^ (St. Louis, MO).

#### Statistics

Statistical analysis was done using GraphPad Prism software (version 11.0.0, Boston, MA). Comparisons of age, body weight, F–I gains, EPSP amplitude, and intrinsic electrical properties of motoneurons were conducted using the nonparametric Kruskal–Wallis test followed by Dunn’s multiple-comparisons post hoc test. The motoneuron firing phenotypes were compared using Chi-square test. Finally, a mixed-effects analysis was used to compare ventral root CAPs across different DR stimulation amplitudes. Statistical significance was denoted as **\***: p<0.05, **: P<0.01, ***: P<0.001, ****: P<0.0001.

## RESULTS

Riluzole is known to acutely suppress motoneuron excitability and glutamate release [4, 26], thus reducing excitotoxicity. In this study, we investigated whether these effects persist following extended riluzole treatment (10 days, in drinking water, Fig. 1). At the end of treatment, the sacral spinal cord was harvested from each animal and used ex vivo to assess motoneuron excitability and synaptic inputs. Importantly, during terminal experiments, the tissue was perfused with a large volume of ACSF (1 L) to washout any remaining riluzole. Hence, electrophysiological measurements were made in absence of the drug, apart from one set of experiments where riluzole was reintroduced ex vivo (see below).

### Effect of chronic riluzole treatment on motoneuron excitability

We evaluated the excitability of motoneurons via their input-output relationship. Using dynamic slowly rising and falling current ramps (Fig. 2A), we established frequency-current (F-I) relationships for both the ascending and descending parts of the ramp. The gain of the F-I relationship in mSOD1 motoneurons increased after riluzole treatment, while WT motoneurons showed no change (Fig. 2B). Moreover, applying comparative voltage ramps in voltage clamp mode showed increased persistent inward currents (PICs) in mSOD1-cRil motoneurons as compared to WT-cRil cells (Table 1). These PICs are major contributors to motoneuron excitability [27-29].

**Table 1:** Effect of chronic riluzole administration on motoneuron electrical properties in WT and mSOD1 mice. Data was analyzed using Kruskal-Wallis test and Dunn’s multiple comparison test. *: p<0.05, **: p<0.01.

|  | WT |  |  | WT-cRil |  |  | mSOD1 |  |  | mSOD1-cRil |  |  |
| --- | --- | --- | --- | --- | --- | --- | --- | --- | --- | --- | --- | --- |
| | Median (IQR) | Mean ± SD | n | Median (IQR) | Mean ± SD | n | Median (IQR) | Mean ± SD | n | Median (IQR) | Mean ± SD | n |
| V<sub>Mem</sub> (mV) | -72.34<br>(-73.55 – -69.15) | -71.29 ± 3.18 | 34 | -69.7<br>(-73.03 – -69.15) | -70.22 ± 4.03 | 42 | -68.2<br>(-71.6 – -65.33) | -68.48 ± 4.79 | 31 | -70.30<br>(-73.63 – -67.75) | -69.16 ± 5.55 | 34 |
| V<sub>th-pulse</sub> (mV) | -52.9<br>(-54.8 – -46.7) | -49.93 ± 7.16 | 31 | **-47.1 \*\*WT**<br>**(-48.7 – -43.4)** | **-46.4 ± 5.02** | 39 | -48.3<br>(-51.33 – -41.9) | -47.18 ± 6 | 28 | -49.8<br>(-51.5 – -46.5) | -48.66 ± 5.19 | 31 |
| V<sub>th-DR stim</sub> (mV) | -50.8<br>(-54.48 – -48.25) | -50.51 ± 4.99 | 30 | -50.8<br>(-52.2 – -48.3) | -50.82 ± 3.63 | 39 | -52.2<br>(-53.9 – -50.3) | -52.17 ± 2.91 | 27 | -50.28<br>(-54.1 – -47.7) | -50.91 ± 3.59 | 33 |
| PIC (nA) | -0.55<br>(-1.32 – -0.29) | -0.86 ± 0.89 | 18 | -0.39<br>(-1.08 – -0.17) | -0.76 ± 1.01 | 16 | -1.24<br>(-1.7 – -0.5) | -1.13 ± 0.73 | 15 | **-1.46 \*WT-cRil**<br>**(-1.87 – -0.66)** | **-1.55 ± 1.17** | 17 |
| Rheobase (nA) | 2.5<br>(1.8 – 3.6) | 2.75 ± 1.23 | 32 | 3<br>(2.4 – 3.95) | 3.22 ± 1.17 | 42 | 3.4<br>(2.3 – 4) | 3.34 ± 1.34 | 29 | 3<br>(1.8 – 4) | 3.03 ± 1.5 | 29 |
| R<sub>peak</sub> (MΩ) | 8.85<br>(7.06 – 11.04) | 9.9 ± 4.03 | 34 | 8.04<br>(7.01 – 9.49) | 8.68 ± 2.59 | 42 | **6.97 \*\*WT**<br>**(5.86 – 8.42)** | **7.06 ± 1.94** | 28 | **8.63 \*mSOD1**<br>**(6.97 – 11.41)** | **9.57 ± 3.55** | 32 |
| R<sub>plateau</sub> (MΩ) | 6.42<br>(5.2 – 8.2) | 7.47 ± 3.512 | 34 | 6.08<br>(5.09 – 7.41) | 6.61 ± 2.46 | 42 | **4.94 \*WT**<br>**(3.68 – 6.55)** | **5.29 ± 1.95** | 28 | **7.16 \*\*mSOD1**<br>**(5.11 – 9.77)** | **7.67 ± 3.39** | 32 |
| Sag ratio | 1.32<br>(1.25 – 1.51) | 1.37 ± 0.16 | 34 | 1.33<br>(1.23 – 1.45) | 1.35 ± 0.15 | 42 | 1.3<br>(1.25 – 1.45) | 1.38 ± 0.21 | 28 | 1.22<br>(1.12 – 1.52) | 1.31 ± 0.22 | 32 |
| Time constant (ms) | 12.19<br>(10.99 – 14.37) | 12.49 ± 3.25 | 33 | 12.94<br>(11.37 – 15.75) | 12.93 ± 3.503 | 39 | 13.67<br>(12.08 – 16.19) | 13.58 ± 2.99 | 22 | 13.14<br>(10.83 – 15.16) | 13.09 ± 2.9 | 20 |
| Electrotonic length | 1.53<br>(1.39 – 1.67) | 1.53 ± 0.19 | 33 | 1.53<br>(1.43 – 1.63) | 1.54 ± 0.17 | 39 | **1.35 \*WT-cRil**<br>**(1.25 – 1.57)** | **1.39 ± 0.22** | 22 | **1.38 \*WT/WT-cRil**<br>**(1.32 – 1.46)** | **1.38 ± 0.14** | 20 |
| G<sub>AHP</sub> (μS) | 0.032<br>(0.017 – 0.042) | 0.038 ± 0.027 | 28 | 0.026<br>(0.02 – 0.034) | 0.03 ± 0.018 | 25 | **0.055 \*WT-cRil**<br>**(0.025 – 0.089)** | **0.062 ± 0.048** | 24 | **0.026 \*mSOD1**<br>**(0.015 – 0.034)** | **0.033 ± 0.033** | 20 |
| G<sub>AHP</sub> / G<sub>in</sub> | 0.32<br>(0.21 – 0.4) | 0.35 ± 0.23 | 28 | 0.23<br>(0.17 – 0.3) | 0.25 ± 0.12 | 25 | 0.33<br>(0.21 – 0.47) | 0.37 ± 0.2 | 24 | 0.23<br>(0.18 – 0.35) | 0.27 ± 0.17 | 20 |
| mAHP amp (mV) | 3.25<br>(1.83 – 4.03) | 3.13 ± 1.49 | 28 | 2.5<br>(1.81 – 3.04) | 2.41 ± 1.11 | 25 | 2.68<br>(1.76 – 3.53) | 2.77 ± 1.45 | 25 | 2.22<br>(1.46 – 2.78) | 2.35 ± 1.21 | 23 |
| mAHP half decay (ms) | 17.72<br>(15.6 – 21.6) | 21.97 ± 11.08 | 28 | 18.8<br>(16.05 – 21.93) | 19.45 ± 4.62 | 25 | 17.6<br>(15.38 – 19.69) | 17.45 ± 2.72 | 25 | 15.45<br>(14.04 – 18.78) | 18.06 ± 8.1 | 24 |
| mAHP V<sub>reversal</sub> (mV) | -85.24<br>(-88.09 – -79.8) | -84.76 ± 7.66 | 28 | -83.78<br>(-88.8 – -77.89) | -84.09 ± 7.7 | 25 | -82.1<br>(-84.24 – -78.7) | -82.13 ± 6.31 | 25 | -82.35<br>(-85.64 – -80.61) | -82.88 ± 4.32 | 23 |

**Fig. 2:**
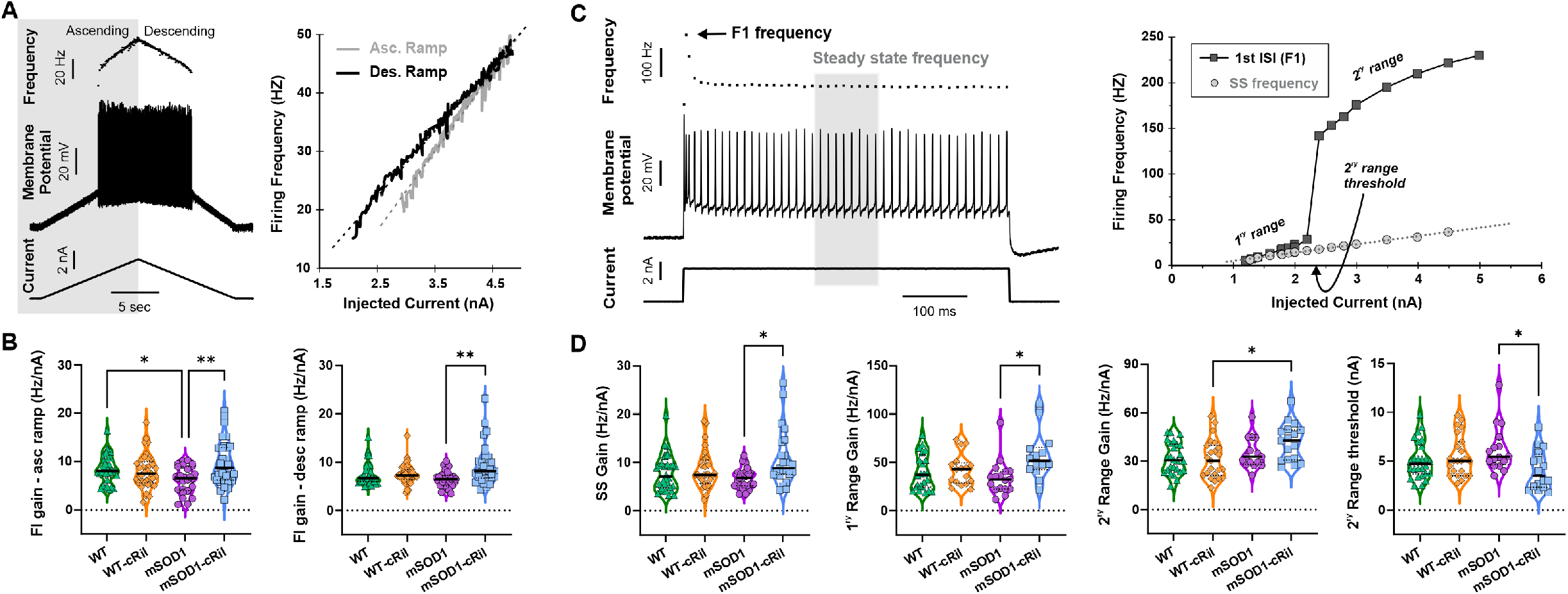
Chronic riluzole treatment selectively increases excitability in mSOD1, but not WT, motoneurons. **A:** Left: Example data showing motoneuron response to intracellular current ramp injection. Right: F-I relationship for the motoneuron shown in (A), depicting the ascending (gray line) and descending (black line) parts of the ramp. **B:** Summary of F-I gain for both the ascending (left) and descending (right) ramp in different experimental groups. The data shows that mSOD1 motoneurons, unlike WT, exhibit increased F-I gain following chronic riluzole treatment. Data was analyzed using Kruskal-Wallis test and Dunn’s multiple comparison test. WT (N= 17 mice, n= 34 MNs), WT-cRil (N= 15, n= 37), mSOD1 (N= 10, n= 26), and mSOD1-cRil mice (N= 15, n= 33). **C:** Left: Example motoneuron response to intracellular current pulse. The instantaneous frequency of the first doublet is marked with an arrow. Steady state firing is measured as the average frequency of the shaded gray area. Right: F-I relationship for the same cell, depicting the first doublet frequency (F1, solid line) and the steady state (SS, dotted line). **D:** Summary of the gain of the F-I relationship of both SS and F1 frequencies (1ry, 2ry, and current threshold for 2^ry^ range). mSOD1 motoneurons exhibit increased F-I gain and reduced current threshold of 2^ry^ range following treatment. Data was analyzed using Kruskal-Wallis test and Dunn’s multiple comparison test. WT (N= 16 mice, n= 32 MNs), WT-cRil (N= 15, n= 31), mSOD1 (N= 10, n= 23), and mSOD1-cRil mice (N= 11, n= 23). *: p<0.05. **: p<0.01.

To determine whether the observed changes in motoneuron excitability were specific to slow depolarizing inputs, we applied a series of fast current pulses with increasing amplitude (see example pulse in Fig. 2C). Motoneurons typically exhibit spike frequency adaptation during the initial part of the pulse (Fig. 2C, left). For each pulse amplitude, we quantified the steady-state firing frequency (SS) and the frequency of the first doublet (F1) and constructed F-I relationships for both metrics (Fig. 2C, right). The SS F-I relationship was linear, whereas F1 displayed a sudden increase in firing frequency after crossing a certain current threshold, prompting its division into primary and secondary ranges. Following extended riluzole treatment, mSOD1-cRil motoneurons exhibited increased SS and F1 gains, along with reduced current threshold for the secondary range (Fig. 2D), indicating enhanced excitability.

Taken together, extended riluzole treatment triggered an increase in motoneuron excitability in mSOD1 mice, while exerting no effect on motoneurons of WT mice.

To assess whether the change in F-I gain altered the overall firing behavior of motoneurons, cells were classified into four distinct types (l - lV) based on their firing behavior during current ramp injection (Fig. 3A, see methods for details). These types are arranged in order of increasing excitability, with type-IV being the most excitable. We then compared the distribution of firing types across experimental groups. WT motoneurons showed no significant change in firing type proportions after riluzole treatment. In contrast, mSOD1 motoneurons exhibited a marked shift toward more excitable types, with the proportion of type-IV cells more than doubling (Fig. 3B).

**Fig. 3:**
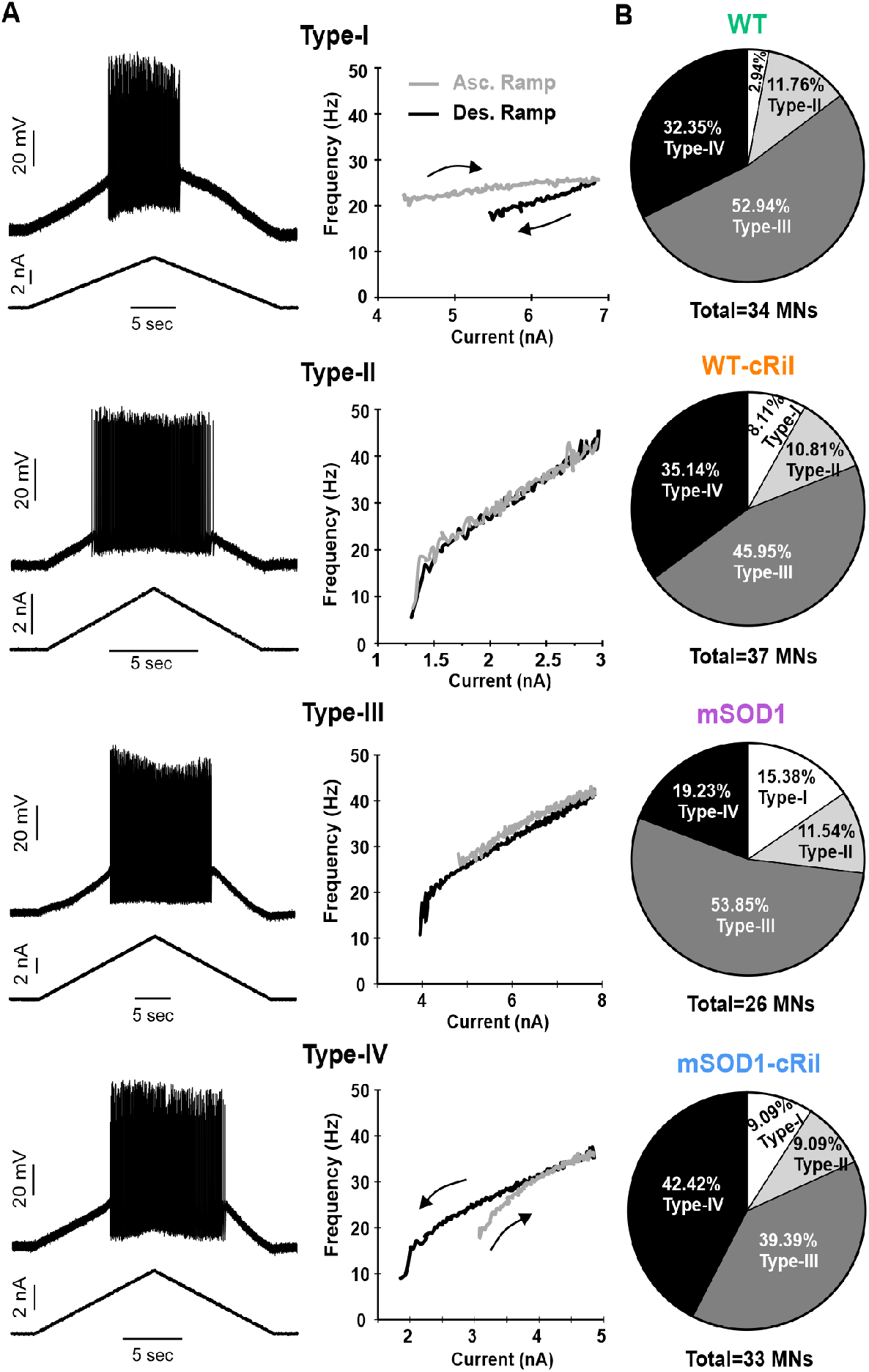
mSOD1 motoneurons exhibit more excitable firing phenotypes following chronic riluzole treatment. **A:** Examples of cellular responses (left) and F-I relationships (right) showing different firing styles (types l - lV) evoked in motoneurons via current ramps. The types are arranged in order of increasing excitability. **B:** Proportions of firing types among different experimental groups. Following riluzole treatment, mSOD1 motoneurons exhibit more excitable firing types (p= 0.0092), while WT motoneurons show no significant change (p = 0.2815). The number of animals and MNs are the same as Figure 2A-B. Data was analyzed using the Chi-square test.

These findings suggest that chronic riluzole treatment induced a homeostatic increase in excitability of mSOD1 motoneurons, while WT motoneurons remain unaffected. This confirms an upregulation in the gain of homeostatic mechanisms in this ALS model.

### Reintroducing riluzole *ex vivo* does not markedly change motoneuron excitability

The increased excitability of mSOD1 motoneurons following chronic riluzole treatment was discovered *ex vivo* after the drug had been washed out. We asked whether the continued presence of riluzole *in vivo* would mask this adaptive increase in excitability. To answer this, we performed a separate set of experiments in which riluzole was acutely reintroduced *ex vivo* (aRil) to spinal cord tissue from WT and mSOD1 mice that had received chronic riluzole treatment (cRil). We used 4 µM of riluzole in these experiments, a clinically relevant concentration [30]. Motoneurons from both WT-cRil and mSOD1-cRil groups showed no significant response to acute riluzole application (Fig. 4).

**Fig. 4:**
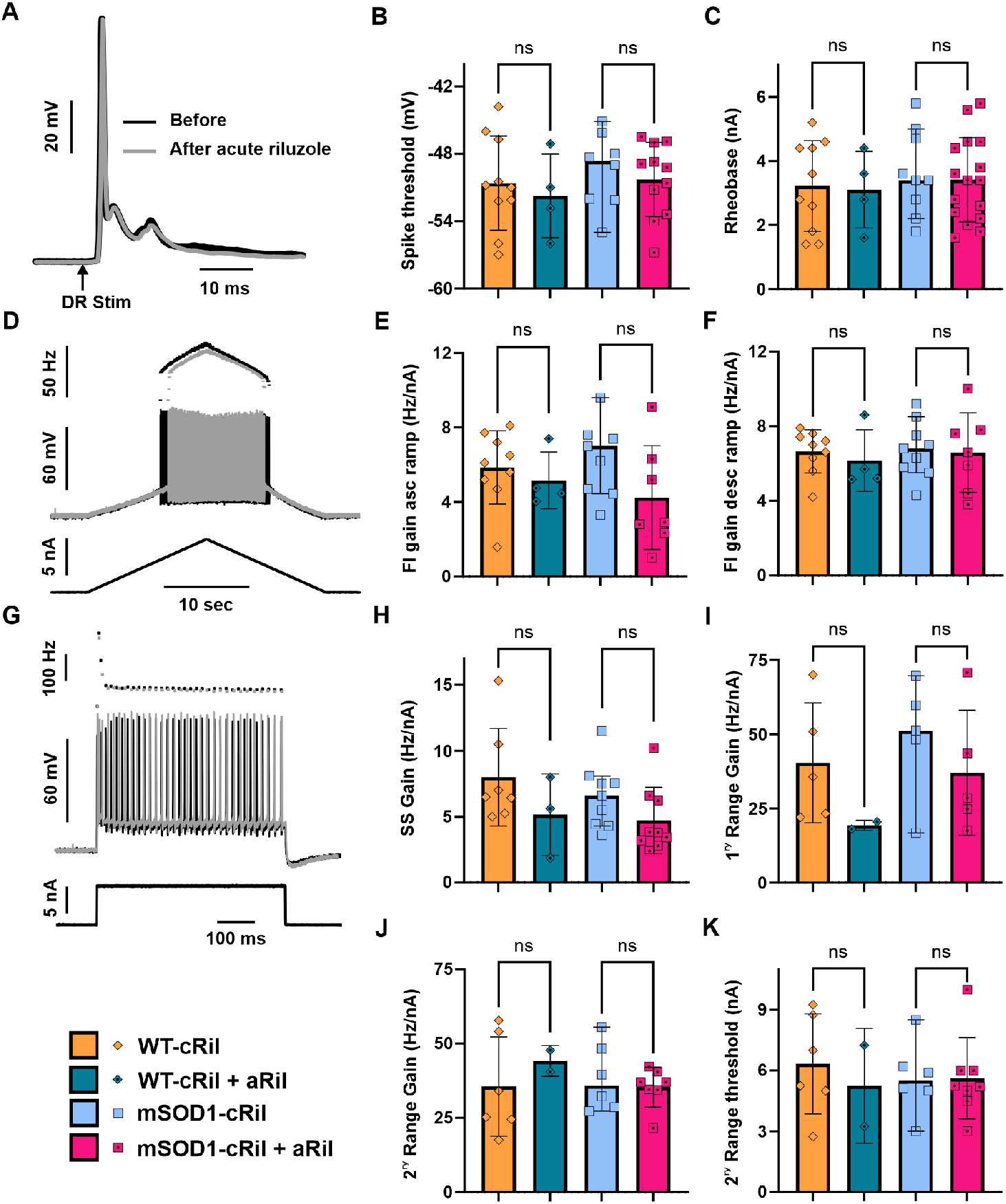
*Ex vivo* riluzole application did not further alter motoneuron excitability in tissue previously exposed to chronic riluzole treatment. Riluzole (4 µM) was applied acutely during terminal experiments (aRil) to spinal cord tissue isolated from WT and mSOD1 mice that had received 10 days of chronic riluzole treatment (cRil). Data obtained before and during acute riluzole application were collected from the same preparations and, in some cases, from the same motoneuron. **A:** Representative intracellular recording from an mSOD1-cRil motoneuron showing an action potential evoked by single-pulse (2×T) dorsal root stimulation before (black) and after (gray) acute riluzole application. **B-C:** Effect of acute riluzole application on the spike voltage threshold and current threshold (rheobase) of motoneurons. **D:** Firing response of the same motoneuron in (A) to ramp current injection before and after riluzole application, illustrating a modest reduction in firing frequency. **E-F:** Summary of the effect of acute riluzole on the gain of the F-I relationship during the ascending (E) and descending (F) phases of the current ramp. **G:** Response of the same motoneuron in (A) to brief depolarizing current pulse before and after acute riluzole application. **H-K:** Effect of acute riluzole on motoneuron firing evoked by current pulses, showing no significant change in steady state F-I gain (H), primary range F1 gain (I), secondary range F1 gain (J), or secondary range current threshold (K). Data was analyzed using Kruskal-Wallis test and Dunn’s multiple comparison test. WT-cRil (N= 4 mice, n= 10 MNs), WT-c+aRil (N= 4, n= 4), mSOD1-cRil (N= 4, n= 9), and mSOD1-c+aRil mice (N= 4, n= 7). ns: non-significant.

In both genotypes, spike threshold measured from action potentials evoked by dorsal root stimulation (Fig. 4A-B), as well as rheobase current (Fig. 4C), were unaffected by acute riluzole. Although riluzole caused a modest reduction in firing frequency in some cells (see Fig. 4D), the overall F-I gain was unchanged (Fig. 4E-F). Similar results were obtained when excitability was evaluated using square pulse current inputs (Fig. 4G-K). Together, these findings indicate that the increased excitability of mSOD1-cRil motoneurons would persist *in vivo*, even in the continued presence of riluzole.

### Effect of chronic riluzole on synaptic inputs to motoneuron

In addition to altering intrinsic neuronal excitability, riluzole can decrease synaptic glutamate release [4, 8]. To evaluate the impact of chronic riluzole on glutamatergic transmission, we recorded ventral root (VR) responses to dorsal root (DR) stimulation ex vivo. Specifically, we measured both the ipsilateral monosynaptic and contralateral polysynaptic VR responses. The activation curves for the ipsilateral monosynaptic reflex showed no significant differences across experimental groups (Fig. 5A-B). In contrast, the contralateral polysynaptic reflexes tended to be larger in treated groups of either genotype compared to their untreated counterparts. However, the increase was significant only for mSOD1-cRil group as compared to untreated mSOD1, and only at low stimulation intensities (Fig. 5C). We also recorded EPSPs in motoneurons following contralateral DR stimulation at 2xT. These EPSPs showed a similar trend toward enhancement in cRil groups but were not statistically significant except for mSOD1-cRil compared to untreated WT (Fig. 5D). This data suggests that chronic riluzole treatment does not induce homeostatic changes in the monosynaptic reflex pathway in either genotype. Nonetheless, riluzole causes an increase in polysynaptic pathway transmission to mSOD1 motoneurons and not WT.

**Fig. 5:**
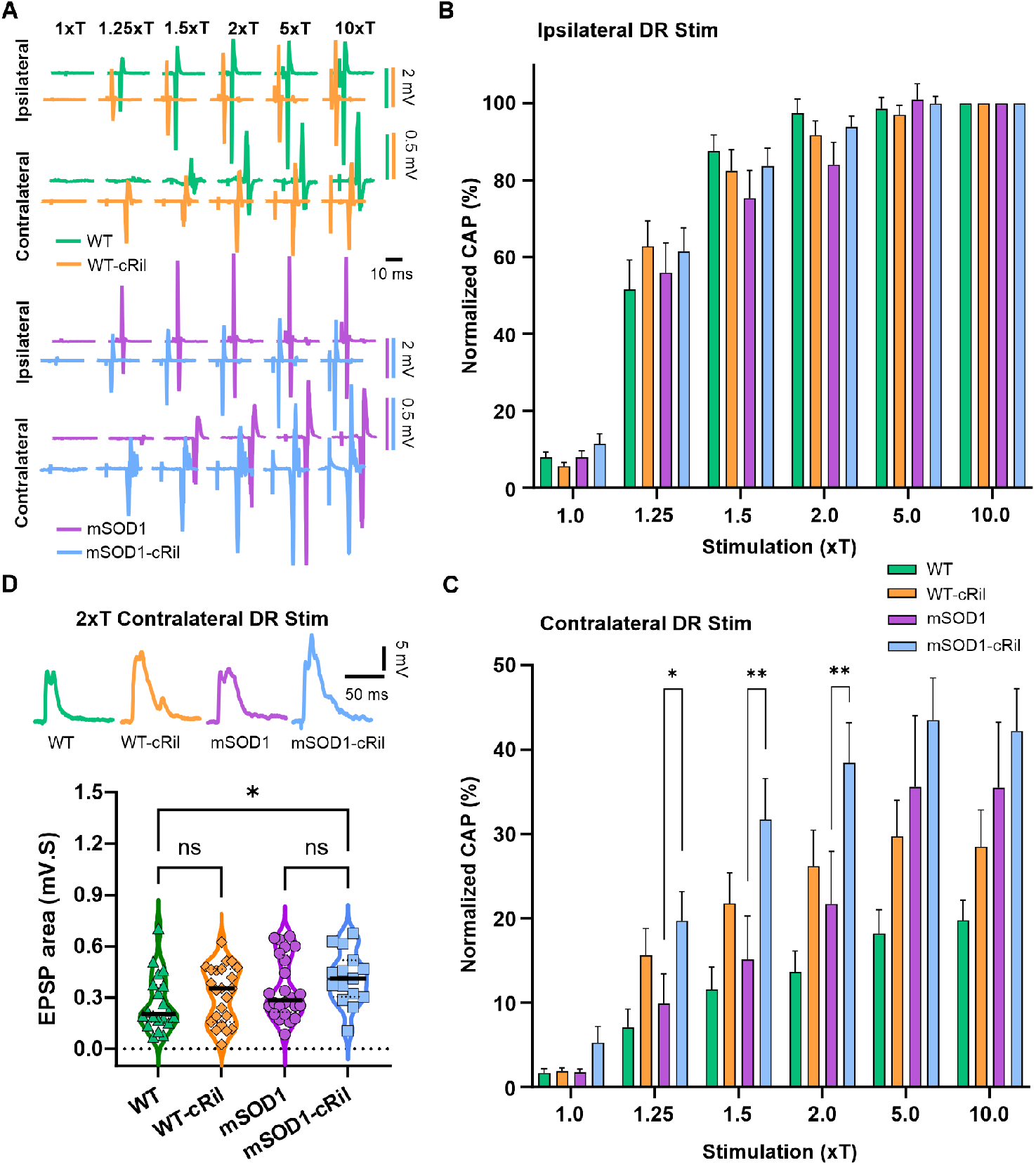
Effect of chronic riluzole on synaptic inputs to motoneurons. **A:** Examples of root reflex data showing ventral root compound action potentials (CAPs) in response to ipsilateral and contralateral dorsal root stimulation at multiple intensities (xThreshold, as indicated at the top. Note: Data from WT-cRil and mSOD-cRil are slightly time-shifted to the left for clarity. **B-C:** Summary of ipsilateral (C) and contralateral (D) CAP responses normalized to the maximum response of the respective root (at 10xT ipsi DR stim). Data is represented as the mean ± SEM. WT (N= 10 mice, n= 19 VRs), WT-cRil (N= 10, n= 19), mSOD1 (N= 10, n= 20), and mSOD1-cRil mice (N= 14, n= 26). **D:** Top: Examples of EPSPs evoked in motoneurons by 2xT contralateral DR stimulation. Bottom: Summary of EPSP measurements (curve area). WT (N= 10 mice, n= 20 MNs), WT-cRil (N= 12, n= 23), mSOD1 (N= 10, n= 24), and mSOD1-cRil mice (N= 8, n= 15). Data in B-C were analyzed using a mixed-effects model. Data in D was analyzed using Kruskal-Wallis test and Dunn’s multiple comparison test. *: p<0.05, **: p<0.01.

### Chronic riluzole modulates passive electrical properties of motoneurons

We next compared the basic electrical properties of recorded motoneurons to investigate whether they contribute to the increased excitability of mSOD1-cRil motoneurons (Fig. 6 and Table 1). Motoneurons of untreated mSOD1 mice had larger input conductance as compared to their WT counterparts (Fig. 6A-B). This is consistent with previous reports showing increased motoneuron size at this stage of the disease [14, 15, 31]. Surprisingly, chronic riluzole treatment caused a decrease in the input conductance of mSOD1 motoneurons and had no effect on WT motoneurons (Fig. 6B-C).

**Fig. 6:**
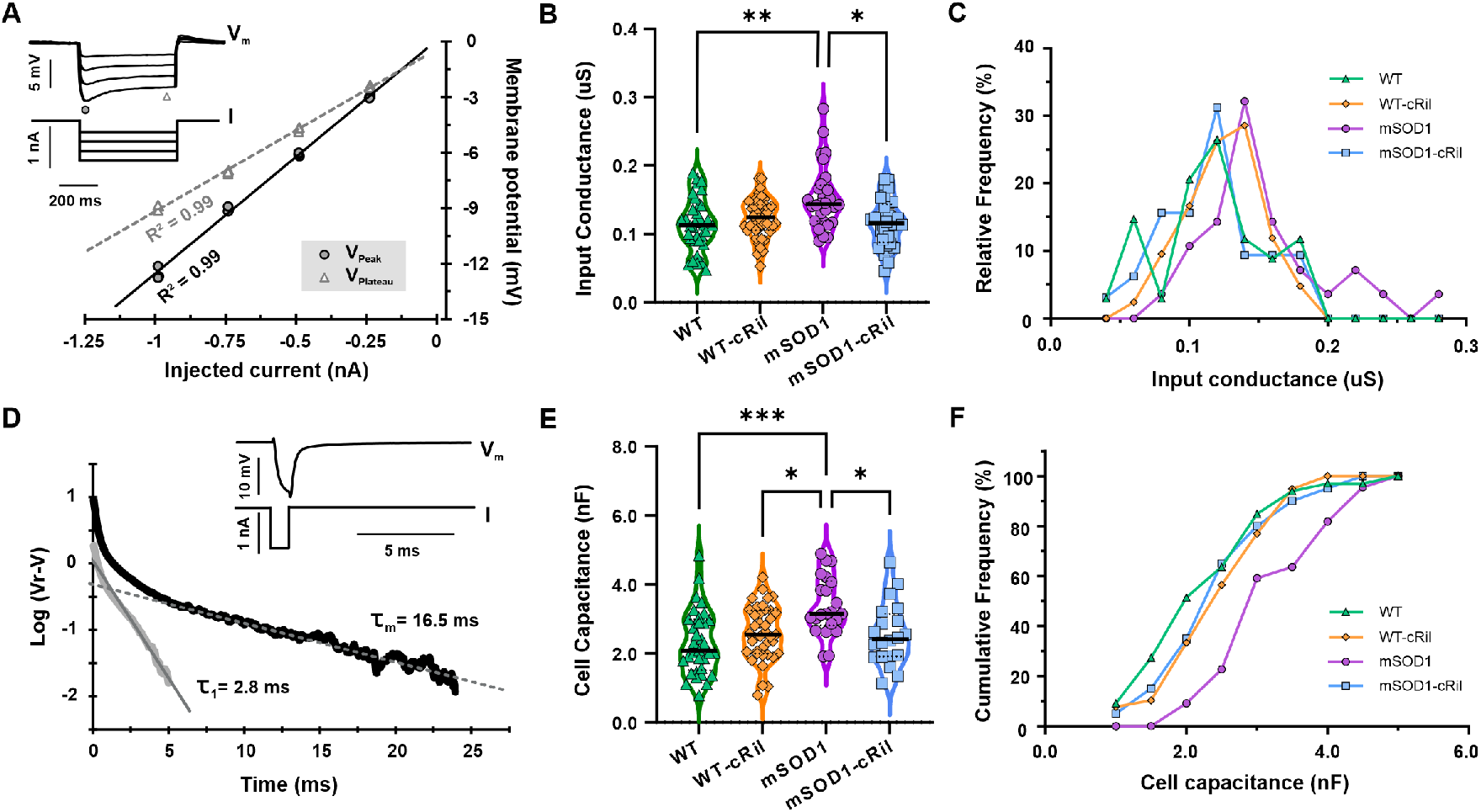
Chronic riluzole administration restores mSOD1 motoneuron size. **A:** Example traces showing overlay of multiple negative current steps used to measure motoneuron input conductance. A current-voltage relationship is established for the peak and plateau voltage values. **B:** Summary of motoneuron input conductance (peak values) for all experimental groups. **C:** Histogram showing the distribution of motoneuron input conductance. WT (N= 17 mice, n= 34 MNs), WT-cRil (N= 15, n= 42), mSOD1 (N= 10, n= 28), and mSOD1-cRil mice (N= 15, n= 32). **D:** Example traces showing average response to a short hyperpolarizing pulse used to estimate motoneuron membrane time constant. A semilogarithmic plot of the voltage response (resting potential subtracted) is established (black), and a straight line is fitted over the linear portion. The slope of this linear fit is used to measure the membrane time constant (τ_m_). The first equalizing time constant (τ_1_) is then estimated by peeling off the extrapolated linear fit from the response (red). **E:** Summary of calculated motoneuron membrane capacitance. **F:** Cumulative histogram showing the distribution of cell capacitance. WT (N= 17 mice, n= 33 MNs), WT-cRil (N= 15, n= 39), mSOD1 (N= 8, n= 22), and mSOD1-cRil mice (N= 9, n= 20). Chronic riluzole treatment shifts input conductance and capacitance of mSOD1 motoneurons to normal range but has no effect on WT motoneurons. Data was analyzed using Kruskal-Wallis test and Dunn’s multiple comparison test. *: p<0.05, **: p<0.01.

The reduction in mSOD1 motoneuron conductance usually reflects a decrease in cell membrane area. However, it could also result from shrinkage of cell volume due to blocking of Na^+^ entry by riluzole. To identify the exact mechanism, we measured membrane time constant (Fig. 6D) and used it to calculate total capacitance (see methods), a more accurate measure of membrane area. Assuming that specific membrane capacitance is constant among motoneurons (about 1 µF/cm^2^), the total cell capacitance gives an accurate estimate of motoneuron surface area [23, 24]. The data showed a similar trend of increased capacitance in untreated mSOD1 vs WT motoneurons and a reduction in mSOD1-cRil motoneuron capacitance following treatment (Fig. 6 E-F). This indicates that chronic riluzole treatment caused a reduction in membrane area/motoneuron size.

There were no major changes in the other electrical properties measured (see Table 1). Nonetheless, the combination of reduced cell size along with increased PICs and unchanged spike threshold and AHP properties (Table 1) can underly the increase in mSOD1-cRil motoneuron excitability following treatment.

Overall, our results demonstrate that chronic riluzole treatment in ALS paradoxically induces an adaptive increase in motoneuron excitability and synaptic excitation. These effects are probably due to increased gain of homeostatic mechanisms in this ALS model [13, 16]. Therefore, our findings challenge the prevailing view of riluzole mechanism of action in ALS. The data instead revealed a previously unrecognized effect of chronic riluzole treatment in ALS, a reduction in motoneuron size. This effect could lower metabolic demand and thereby contribute to prolonged cell survival.

## DISCUSSION

The therapeutic benefits of riluzole in ALS appear to diminish after several months of treatment [17]. Using intracellular recordings, we investigated the long-term effects of riluzole on adult motoneurons of WT and presymptomatic mSOD1 mice. The data showed that chronic treatment increased excitability and excitatory inputs of mSOD1 motoneurons, which may contribute to the limited clinical efficacy of the drug. In addition, we also found that riluzole can normalize motoneuron size, potentially reducing cellular metabolic demands.

### Effects of riluzole on neuronal excitability

Riluzole is known to inhibit voltage-gated Na^+^ channels, preferentially when the channel is in the inactivated state, which is more prevalent in depolarized or actively firing neurons [32, 33]. As a result, riluzole reduces the ability of active neurons to generate action potentials, thereby decreasing both presynaptic and postsynaptic excitability. Reduced presynaptic excitability leads to diminished release of excitatory neurotransmitters, among others, from synaptic terminals [8, 34]. In parallel, it decreases postsynaptic responsiveness to depolarizing inputs. Also, riluzole has been shown to suppress the persistent inward Na^+^ current (Na^+^ PIC) [4, 7, 35], a conductance that plays a key role in spike initiation and the maintenance of repetitive firing in spinal motoneurons [6]. Collectively, these actions reduce motoneuron excitability and could, in turn, engage compensatory homeostatic mechanisms.

### Dysregulated homeostasis in ALS motoneurons

Temporal analyses of multiple biological parameters across disease progression in the mSOD1 mouse have revealed slow oscillatory patterns, with alternating decreases and increases at different disease stages [12]. These parameters include mitochondrial function, Ca^2+^ regulation, and oxidative stress markers. Subsequent studies have reported similar oscillatory trends in motoneuron soma size and intrinsic excitability [14, 15]. Such dynamics can arise in biological systems, as in engineered ones, when feedback regulation operates with high gain and sufficiently delayed responses. Accordingly, it has been proposed that these oscillations in ALS reflect an abnormally increased gain of cellular feedback mechanisms, often described as “hypervigilant” homeostasis [13]. Within this framework, mSOD1 motoneurons respond excessively to perturbations, leading to overcompensation that initiates successive cycles of correction; thereby creates the observed oscillations. In the present study, we further examined this idea by treating mSOD1 mice with therapeutic levels of riluzole for an extended period, then measured motoneuron excitability.

### Riluzole and hyperactive homeostasis in ALS

The neuroprotective effects of riluzole in ALS have generally been attributed to its ability to reduce motoneuron excitability and glutamate release [10]. Our results show that after 10 days of riluzole administration in vivo, motoneurons of mSOD1 mice became more excitable. This was reflected by an increased F–I gain and a shift toward more excitable firing phenotypes. The absence of similar effects in WT animals suggests that the suppressive actions of riluzole indeed triggered an exaggerated homeostatic response in this ALS model, resulting in a compensatory increase in motoneuron excitability and polysynaptic excitatory inputs.

Notably, when riluzole was reintroduced ex vivo, neither WT nor mSOD1 tissue that had been chronically exposed to riluzole showed significant additional changes. This suggests that the adaptive increase in motoneuron excitability in mSOD1 animals may not be fully counteracted by the continued presence of riluzole in vivo, potentially reducing the drug effectiveness over time. Consistent with our data, longitudinal studies in ALS patients show that effects of riluzole on excitability fade over few weeks of continued treatment [17]. Yet, it remains unclear whether riluzole can produce these hyper-compensatory adaptations at all stages of the disease, or whether they are limited to specific stages. Similarly, homeostatic responses to riluzole are yet to be tested in other models of ALS. Addressing these questions may help explain the variability in reported therapeutic outcomes of riluzole, where some studies have found little or no benefit [36, 37]. Importantly, our data indicates that the increase in excitability stems from a slight increase in PICs combined with significant reduction in cell size.

### Effect of riluzole on mSOD1 motoneuron size

Multiple studies have consistently reported that presymptomatic stages in the mSOD1 mouse model are characterized by enlarged motoneurons [15, 16, 31, 38]. This increase in motoneuron size during early adulthood is probably a compensatory response to the heightened motoneuron excitability observed during embryonic and early postnatal development [39-41]. A larger cell size requires greater excitatory input to reach the firing threshold, which may help stabilize neuronal activity. However, increased size also elevates metabolic demand. Indeed, motoneurons exhibit size-dependent vulnerability in ALS, with the largest motoneurons degenerating first [42, 43]. Although the precise reasons for this remain unclear, the greater metabolic demand of large cells is thought to contribute to their increased susceptibility.

Our data shows that prolonged riluzole treatment in mSOD1 mice caused reduction in electrical indicators of total membrane area, suggesting a decrease in overall motoneuron size. This change possibly reflects a homeostatic adjustment to the riluzole-induced initial suppression of network excitability, as it was not observed in cultured motoneurons [44]. The coupled changes in motoneuron size and excitability in ALS have been reported in previous studies, where early postnatal interventions that reduced motoneuron size resulted in adaptive decreases in excitability [16]. Therefore, the reduction in cell size does not seem to be a direct effect of riluzole and is less likely to occur when riluzole is used for neurological conditions that do not involve heightened homeostatic activity.

Given that riluzole seems to lose its anticipated effects on excitability and synaptic inputs during continued treatment, this previously-unrecognized reduction in motoneuron size may represent a crucial therapeutic benefit for the drug in ALS.

### Clinical implications and limitations

Despite being backed by clinical observations, hypervigilant homeostasis has been primarily studied in the mSOD1 mouse model. Thus, further investigation in other animal models of ALS and in ALS patients are needed. In addition, our recordings were obtained ex vivo and in absence of neuromodulatory inputs, such as descending brainstem projections, which are crucial for motoneuron excitability [28, 45]. However, neuromodulators are expected to amplify the differences in excitability we found.

The timing and duration of riluzole treatment may be critical to avoid progressive homeostatic antagonism of drug effects. Early or intermittent treatment strategies may prove more effective than continuous administration. Additionally, the normalization of motoneuron size by riluzole might prove to be a major therapeutic benefit and thus warrants further investigation.

## RESOURCE AVAILABILITY

Data reported in this paper will be made available upon request. Requests for information should be directed to the lead contact, CJ Heckman.

## ACKNOWLEDGMENTS

The authors would like to thank Dr. Matthieu Chardon for his insightful inputs about data analysis. This study was supported by the National Institute of Health (NIH NINDS NS110953) and the Les Turner ALS Foundation.

## AUTHOR CONTRIBUTIONS

CJH and AAM conceived the study and designed the experiments. AAM designed and implemented hardware and software tools for the experiments. AAM and BSH participated in animal care and drug administration. AAM performed the electrophysiology experiments, collected the data, performed the analyses, generated the figures, and wrote the initial draft of the manuscript. All authors contributed to manuscript editing and approved the final version. CJH supervised the project and participated in data interpretation.

